# A comprehensive neural simulation of slow-wave sleep and highly responsive wakefulness dynamics

**DOI:** 10.1101/2021.08.31.458365

**Authors:** Jennifer S. Goldman, Lionel Kusch, Bahar Hazal Yalçinkaya, Damien Depannemaecker, Trang-Anh E. Nghiem, Viktor Jirsa, Alain Destexhe

**Affiliations:** Paris Saclay University, Institute of Neuroscience, CNRS, Gif-sur-Yvette, France; Institut de Neurosciences des Systémes, Aix-Marseille University, INSERM, Marseille, France; Laboratoire de Physique de l’Ecole Normale Supérieure, ENS, Université PSL, CNRS, Sorbonne Université, Université de Paris, Paris

**Keywords:** Neural simulation, biophysics, mean-field model, spontaneous activity, evoked responses, wake, synchronous, slow-wave sleep, human brain

## Abstract

Hallmarks of neural dynamics during healthy human brain states span spatial scales from neuromodulators acting on microscopic ion channels to macroscopic changes in communication between brain regions. Developing a scale-integrated understanding of neural dynamics has therefore remained challenging. Here, we perform the integration across scales using mean-field modeling of Adaptive Exponential (AdEx) neurons, explicitly incorporating intrinsic properties of excitatory and inhibitory neurons. We report that when AdEx mean-field neural populations are connected via structural tracts defined by the human connectome, macroscopic dynamics resembling human brain activity emerge. Importantly, the model can qualitatively and quantitatively account for properties of empirical spontaneous and stimulus-evoked dynamics in the space, time, phase, and frequency domains. Remarkably, the model also reproduces brain-wide enhanced responsiveness and capacity to encode information particularly during wake-like states, as quantified using the perturbational complexity index. The model was run using The Virtual Brain (TVB) simulator, and is open-access in EBRAINS. This approach not only provides a scale-integrated understanding of brain states and their underlying mechanisms, but also open access tools to investigate brain responsiveness, toward producing a more unified, formal understanding of experimental data from conscious and unconscious states, as well as their associated pathologies.

## 1 Introduction

Brain activity is marked by complex spontaneous dynamics, particularly during conscious states when the brain is most responsive to stimuli. Though changes in spontaneous and evoked dynamics have been unambiguously empirically observed in relation to changes in brain state, their multi-scale nature has notoriously occluded a formal understanding.

Spanning from macroscopic dynamics supporting communication between brain regions to microscopic, molecular mechanisms modulating ion channels, hallmarks of consciousness have been observed across spatial scales. At the whole-brain level, conscious states are marked by complex spontaneous collective neural dynamics [1, 2] and more sustained, reliable, and complex responses to stimuli [3, 4, 5, 6]. At the microscopic level, neuromodulation is enhanced in conscious, active states, leading to microscopic changes in cellular kinetics [7]. Yet, a challenging multi-scale problem still resides in comprehending how changes in the complexity of global spontaneous dynamics and whole brain responsiveness may specifically relate to microscopic neuromodulatory processes to enable neural coding during active states. Here, using mean-field models of conductance-based, Adaptive Exponential (AdEx) integrate-and-fire neurons with spike-frequency adaptation developed recently [8, 9, 10], constrained by human anatomy and empirically informed by local circuit parameters, we report the natural emergence of global dynamics mimicking different human brain states.

To connect microscales (neurons) to macroscales (whole brain), this work relies on previous advances at mesoscales (neural populations). The first step was modeling biologically-realistic activity states in networks of spiking neurons. Based on experimental recordings, we used the Adaptive Exponential (AdEx) integrate and fire model to simulate two main cell types identifiable in extracellular recordings of human brain [11]: regular-spiking (RS) excitatory and fast-spiking (FS) inhibitory cells. AdEx networks were constrained by biophysical representations of synaptic conductances, which allowed the model to be compared to conductance measurements done in awake animals [8] (for experiments, see [12, 13]). In such configurations, AdEx networks reproduce states observed *in vivo* [14, 8, 15, 16, 17], notably asynchronous irregular (AI) states found experimentally in awake subjects, and synchronous slow waves as in deep sleep [13, 18, 19]. From AdEx networks, mean-field models were derived to take into account second order statistics of AdEx networks interacting through conductance-based synapses. We used a Master Equation formalism [20], modified to include adaptation [9].

In this manuscript, we present evidence that mean-field descriptions of biophysically informed estimates of neuron networks produce macroscopic dynamics capturing essential characteristics of human wake and sleep states - due to variation in spike-frequency adaptation - when coupled by the human connectome with tract-specific delays. First, we show that simulated microscopic changes in membrane currents directly lead to the emergence of globally asynchronous versus synchronous dynamics exhibiting distinct signatures in the frequency domain, as well as changes in inter-regional correlation structure and phase-locking, mimicking aspects of spontaneous human brain dynamics. Further, we report enhanced brain-scale responsiveness to stimulation in simulations of asynchronous, fluctuation-driven compared to synchronous, phase-locked regimes, consistent with empirical data from conscious versus unconscious brain states. Together, the data suggest that the TVB-AdEx model represents a scale-integrated neuroinformatics framework capable of recapitulating known features associated with human brain states as well as elucidating relationships between space-time scales in brain activity. Due to its reliance on anatomical data non-invasively available from humans, this model may further facilitate subject-specific modeling of human brain states in health and disease, including restful and active waking states, as well as sleep, anaesthesia, and coma to aid future advances in personalized medicine.

## 2 Results

We begin by showing essential properties of the components forming the TVB-AdEx model. Next, we describe the integration of AdEx mean-fields into The Virutal Brain (TVB) simulator of EBRAINS, making the models and analyses openly available to facilitate replication and extension of the results. The results presented here indicate that the TVB-AdEx whole human brain model captures fundamental aspects of synchronous and asynchronous brain states, both spontaneously and in response to perturbation.

### 2.1 Components of TVB-AdEx models

The first component of the TVB-AdEx model is at the cellular level, and consists of networks of integrate-and-fire adaptive exponential (AdEx) neurons. As shown in previous studies [14, 8, 9], networks of AdEx neurons with adaptation can display asynchronous, irregular (AI) states, as well as synchronous, regular slow-wave dynamics that alternate between periods of high activity (Up) and periods of near silence (Down). The necessary mechanistic ingredients needed to obtain both dynamical regimes include leak conductance and conductance-based synaptic inputs. Each neuron’s input is comprised by the firing rates of synaptically connected neurons, weighted by synaptic strengths, as well as stochastic noise (hereafter called “drive”; see Materials and Methods), related biologically to miniature postsynaptic currents. AdEx neurons have the ability to integrate synaptic inputs and fire action potentials, followed by a refractory period [21]. AdEx networks with conductance-based synapses can capture features offered by more detailed and computationally expensive models, including AI states and slow-wave dynamics. Figure 1 shows an example of such AI states (Fig. 1A) and Up-Down dynamics (Fig. 1B) simulated by the same AdEx network, changing only the level of spike-frequency adaptation current (parameter *b* in the equations, see Material and Methods). In AI states, the firing of individual units remains irregular, but sustained (Fig. 1A), whereas in slow-wave states the dynamics alternate between depolarized Up states with asynchronous dynamics and hyperpolarized Down states of near silence (Fig. 1B). As such, changes in spike-frequency adaptation lead to differences in cellular kinetics between sleep and wake states. Biologically, inhibition of spike-frequency adaptation can be induced by enhanced concentrations of neuromodulators such as acetylcholine during active, conscious brain states that tends to close K^+^ leak channels, resulting in sustained depolarization of neurons [7] which promotes the emergence of asynchronous, irregular (AI) action potential firing and fluctuation-driven regimes associated with waking states. In contrast, low levels of neuromodulation during unconscious brain states leave leak K^+^ channels open, leading to waves of synchronous depolarisation and hyperpolarization due to the buildup and decay of spike-frequency adaptation, accounting for the emergence of slow-wave dynamics as observed in previous modeling work [16, 17].

**Figure 1.**
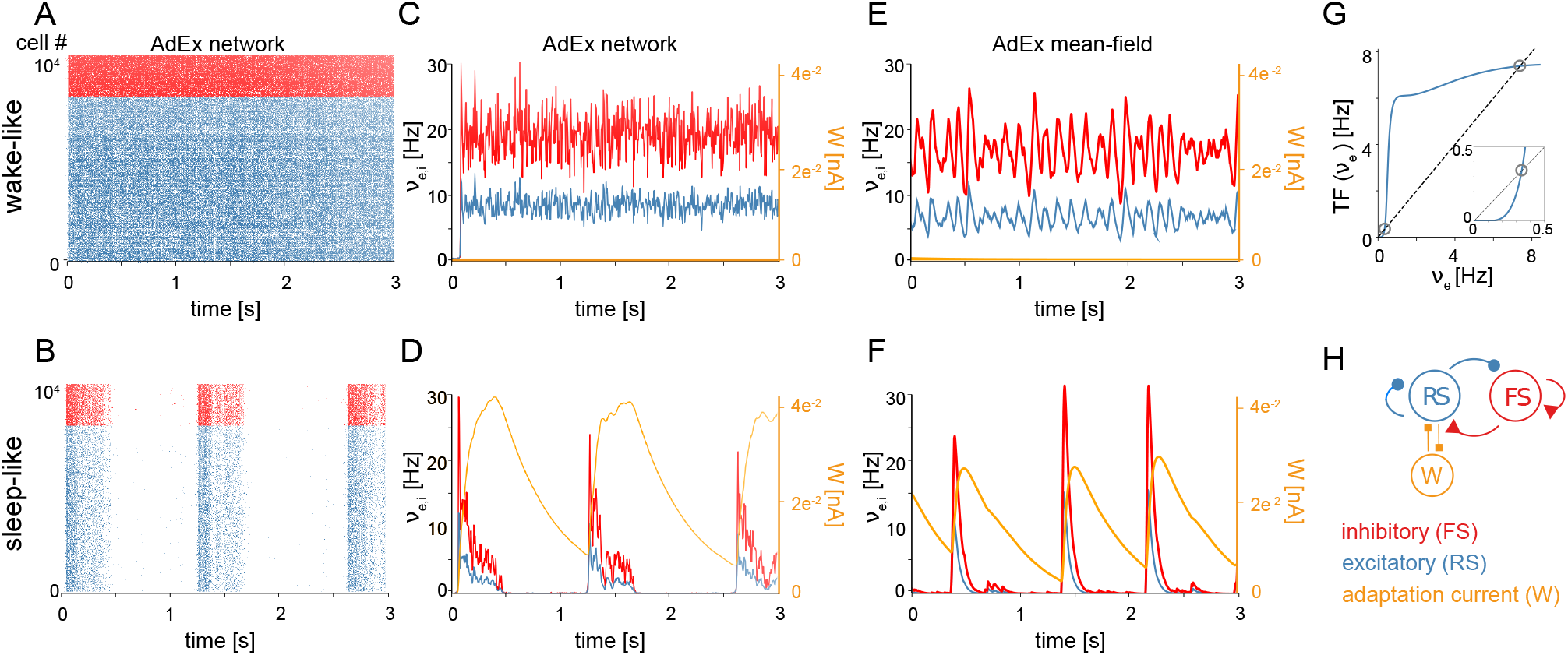
Asynchronous and synchronous dynamics produced by networks of microscopic AdEx neurons and their mesoscopic approximations. Raster plots (A-B) and mean firing rates (C-F) from networks comprised of excitatory RS (blue) and inhibitory FS (red) AdEx neurons displaying asynchronous (A) versus synchronous states (B) as in [9, 10]. The two simulated states, mimicking wake and sleep neural dynamics, differ only in the spike-frequency adaptation current *b_e_* provided to the model (*b_e_* = 0*pA* in the asynchronous state and 60*pA* in the synchronous state), known to be regulated by neuromodulation *in vivo* [7]. C-D. Time variation of mean firing rates (*ν_e,i_*) and adaptation current (*W_e_*) corresponding to networks shown in A-B. Asynchronous (E) and synchronous (F) firing rate dynamics produced using a mean-field model of AdEx networks implemented in The Virtual Brain (TVB). (G) Input-output firing rate relations are given by the mean-field model transfer function (TF). Mean output firing rates of excitatory (blue) neurons as a function of mean excitatory input. The dashed black trace is the identity line. Fixed points of the system (grey circles) occur where the input-output relation intersects with the identity at the positions marked by circles (see Methods for equations). Note that two fixed points are apparent, one at high firing rates and one at low firing rates. The inset is an enlargement of the low-input, low-output region, highlighting the presence of the low-firing fixed point. During asynchronous, wake-like states, firing rates fluctuate around the higher fixed point. During sleep-like states, spike-frequency adaptation builds up as excitatory neurons fire at the high-rate fixed point, eventually destabilizing the high-rate fixed point and causing the system to transition to the near-zero rate fixed point. While the neurons are near-silent, adaptation decays through time, allowing noise fluctuations to entrain a transition back to the high-rate fixed point. (H) Schematic of the simulated network.

The second component of the TVB-AdEx model is a mean-field equation derived from spiking-neuron network simulations, capturing the typical dynamics of a neuron in response to inputs and hence able to describe the mean behaviour of a neuronal population, [8, 9] using a Master-Equation formalism [20]. This formalism allows one to derive mean fields from conductance-based integrate-and-fire models. It has been shown that - using numerical fits of the transfer function [22], an analytical expression for the relationship between a neuron’s input and output rates - one can describe complex neuronal models, such as AdEx neurons, and even Hodgkin-Huxley type biophysical models [23]. In Fig. 1C-D, excitatory and inhibitory firing rates are compared between mean-field simulations using the Master Equation formalism and spiking neural network simulations (time binned population spike counts divided by time bin length T = 0.5 ms). The average adaptation value is also shown for this network (Fig. 1C and D, orange curves). These population variables are suitably captured by the mean-field model including adaptation [9, 10]. This mean-field model can exhibit both AI (Fig. 1E) and Up-Down dynamics (Fig. 1F). Like in the AdEx spiking network model, the transition between two states can be obtained by changing the adaptation parameter called *b* in both cases [9]. With no adaptation, the dynamics are fluctuation-driven around a fixed point exhibiting nonzero firing rates. With adaptation, as the neurons self-inhibit due to adaptation, the nonzero rate fixed point is progressively destabilized by adaptation buildup, driving the dynamics back to the near zero firing rate fixed point until the adaptation wears off and noise drives the system back around the higher-rate fixed point. Thus, with adaptation, the system displays noise-driven alternation between the two fixed-points to generate slow waves (Fig. 1G). These regimes are achieved using the mean-field model, which describes excitatory (RS) and inhibitory (FS) population firing rates as well as the mean adaptation level of excitatory populations (Fig. 1H).

### 2.2 Integration of AdEx mean-field models in TVB

We have used the simulation engines of the Human Brain Project’s (HBP’s) EBRAINS neuroscience research infrastructure (https://ebrains.eu and The Virtual Brain https://ebrains.eu/service/the-virtual-brain) to make access as wide as possible. Replication of the TVB-AdEx findings can be done here, with a free EBRAINS account, and users can clone the repositories to further test or extend the present capacities. The models can also be downloaded from Github at https://gitlab.ebrains.eu/kancourt/tvb-adex-showcase3-git to run locally.

Here, the connection of mean-field models was defined by human tractography data (https://zenodo.org/record/4263723, Berlin subjects/QL_20120814) from the Berlin empirical data processing pipeline [24] (Fig. 2A). A parcellation of 68 regions was used to place localized mean-field models, with long-range excitatory connections (Fig. 2B) and delays (Fig. 2C) defined by tract length and weight estimates in human diffusion magnetic resonance imaging (dMRI) data [25]. Now it becomes possible to simulate brain-scale networks using AdEx-based mean-field models in TVB, hence the name “TVB-AdEx” model.

**Figure 2.**
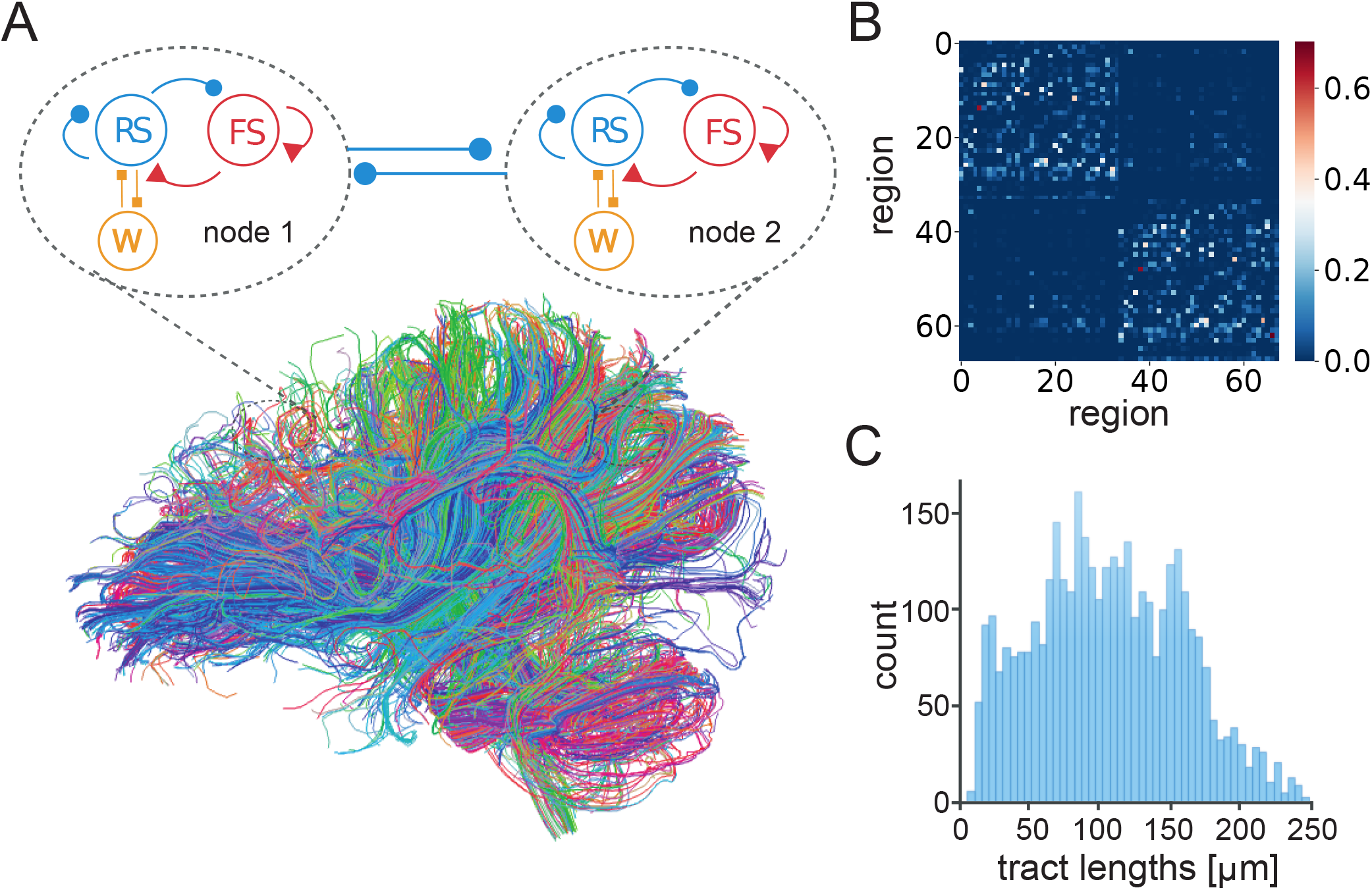
Connection of AdEx mean-field models in The Virtual Brain. (A) Each mean-field model consists of two populations, excitatory RS (blue) and inhibitory FS (red) neurons (as in Fig.1H), taking into account spike-frequency adaptation for excitatory neurons (*W*, orange). Mean-field models represent the mesoscopic scale, here comprising each of 68 defined regions of cerebral cortex. Brain regions are connected by excitatory tracts (thick blue lines) following structural connectomes [24]. (B) Number of fibers connecting brain regions in tractography data, divided by the sum of the gray matter volume of regions in anatomical MRI, is used to define connectivity weights between nodes. (C) The distribution of tract lengths in tractography data informs delays between TVB-AdEx model nodes.

### 2.3 Spontaneous dynamics of large-scale networks

Having coupled AdEx mean-field models that capture average microscopic characteristics of neural activity, we sought to ascertain if hallmarks of brain-scale (macroscopic) spontaneous activity resembling human brain states were reproduced, as well as whether increases in adaptation strength could account for transitions between wake-like and sleep-like macroscopic dynamics. Characterizing temporal hallmarks of simulated neural activity (fig. 3), we find that asynchronous, wake-like dynamics across nodes are recovered in the absence (*b* = 0*pA*, 3A), but not in the presence (*b* = 60*pA*, fig. 3B) of adaptation. Power spectral analysis reveals a peak around 1*Hz* in the high-adaptation condition (fig. 3D) consistent with empirical reports from deeply sleeping individuals, whereas the power spectrum in the low-adaptation condition is more scale-invariant, without characteristic peaks, consistent with irregular dynamics empirically seen during wakefulness (fig. 3C). Therefore, changes in a simulated microscopic process (spike-frequency adaption) can qualitatively reproduce temporal features of macroscopic brain states, with irregular low-adaptation regimes resembling waking states and high-adaptation regimes reminiscent of slow-wave sleep.

**Figure 3.**
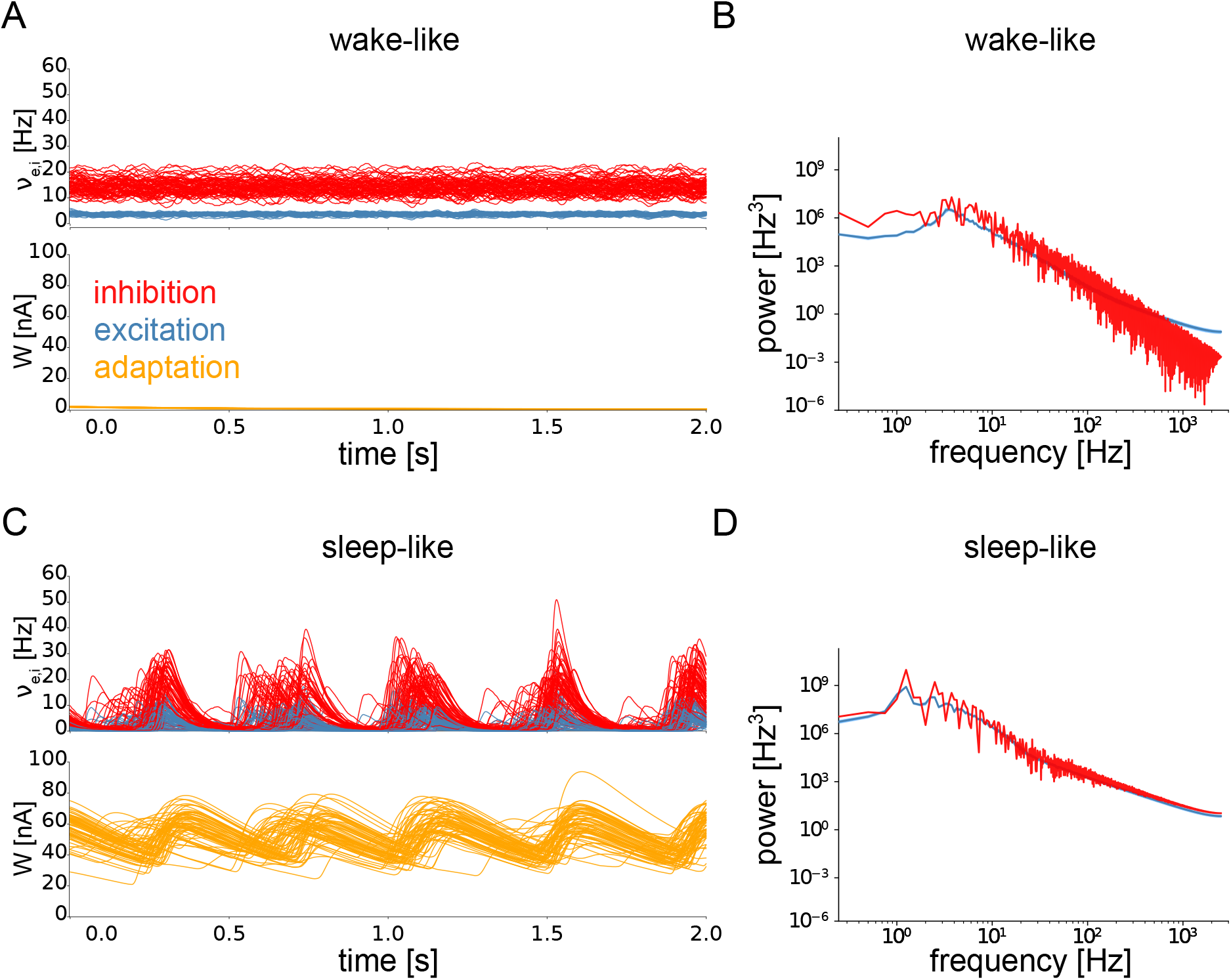
Brain-scale simulations of connected AdEx mean-field models produce activity mimicking wake- and sleep-like states. Time variation of firing rates (*ν_e,i_*, top) and adaptation currents (*W_e_*, bottom) in simulated wake- (A) and sleep-like (B) states for each of the model nodes representing 68 brain regions. When adaptation (*b_e_*) equals 0 pA, the activity of model nodes is asynchronous (A), whereas the inclusion of adaptation (*b_e_* = 60*pA*) leads to the emergence of synchrony between brain regions (B). Fourier power spectra of signals produced by the TVB-AdEx in synchronous (sleep-like, C) and asynchronous (wake-like, D) states. Note that maximal power in the sleep-like condition falls in the delta range (1-4 Hz) while more power is observed at high frequencies in the wake-like state.

Increasing adaptation can also tune the spatial correlation structure of neural activity across brain states. Indeed, as shown in fig. 4, Pearson correlations across nodes are enhanced in the presence of adaptation, consistent with asynchronous dynamics seen during wakefulness versus synchronous slow waves seen during deep sleep (fig. 4A,C). In addition, increased adaptation strength also causes the emergence of significantly larger correlations between inhibitory than excitatory firing rates across nodes during sleep-like dynamics (fig. 4B,D). This reveals that microscopic variation in adaptation strength alone can successfully account for empirical reports of increased correlations between inhibitory neurons across long distances and even different cortical regions for inhibitory but not excitatory neurons [11, 26, 27]. This is due to different effects of adaptation on excitatory regular-spiking neurons and inhibitory fast-spiking neurons, key to reproducing empirical dynamics in unconscious states [16, 17]. Moreover, the Phase Lag Index (PLI) is increased during sleep-like dynamics (4E,G), suggesting systematic phase relations between nodes consistent with travelling slow waves empirically observed during spontaneous unconscious dynamics [18, 19]. Such phase relations, evidenced by a significantly larger PLI, are more pronounced for inhibitory than excitatory neurons in sleep-like dynamics, reminiscent of the key role of inhibitory neurons in organizing the emergence of synchronous dynamics during sleep [28].

**Figure 4.**
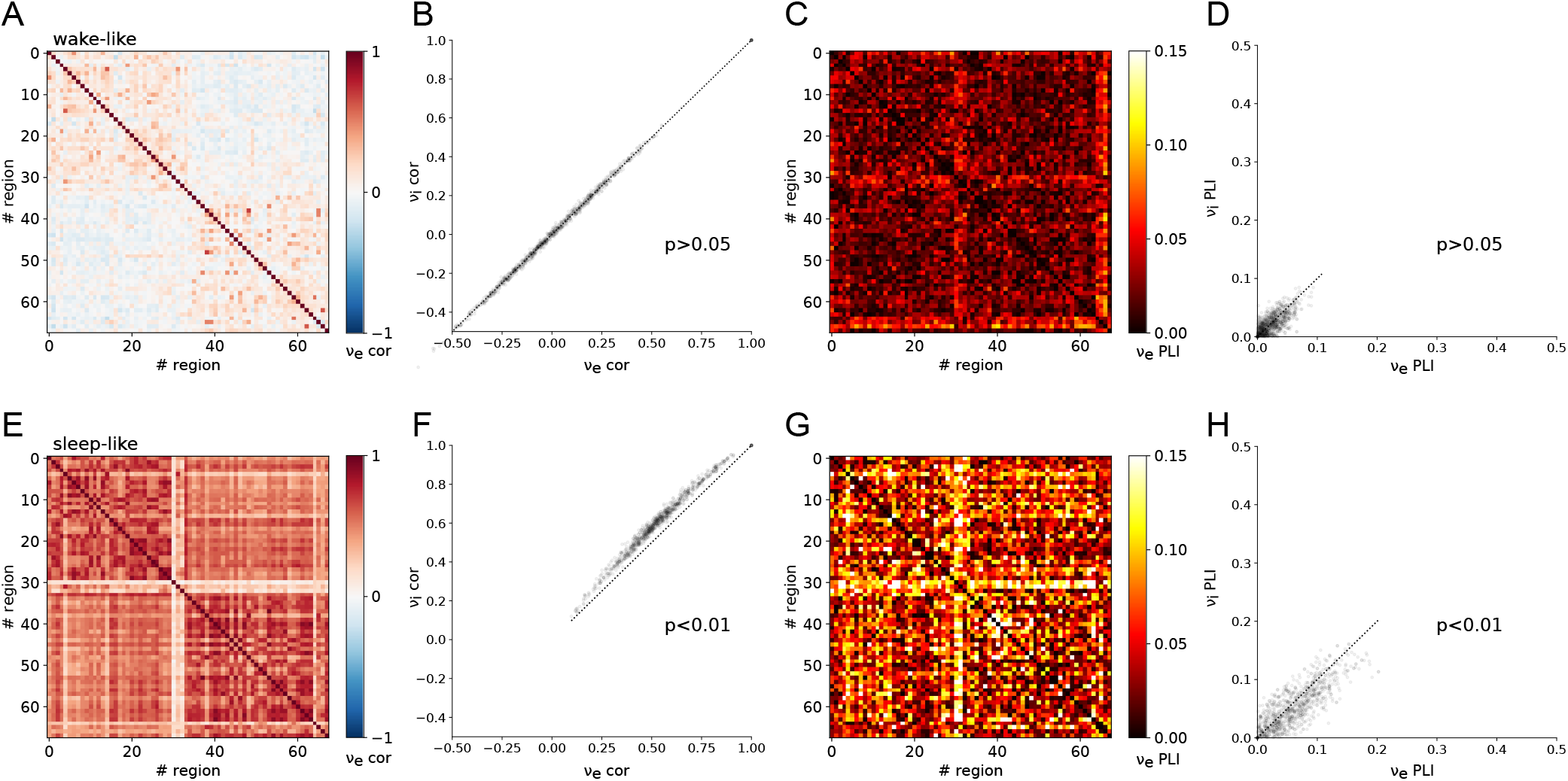
Emergence of enhanced spontaneous synchrony between brain regions in sleep-like simulations. Pearson correlation (A-D) and Phase-Lag Index (PLI) (E-H) in wake-like (A-B, E-F) and deep sleep-like (C-D, G-H) states, heatmaps for correlationsbetween brain regions in terms of excitatory firing rates (A-C, E-G) and scatter plots of inhibitory vs excitatory firing rate correlations (B-D, F-H) where the dotted trace is the identity line. Correlations are overall increased across regions in sleep-like states (C) as compared to wake-like states (A), consistent with increased synchrony across brain regions in empirical brain imaging studies (M/EEG). Correlations across nodes are significantly larger between inhibitory firing rates than between excitatory firing rates in sleep-like dynamics (D; Independent Student’s T-test, *t* = −10, *p* = 1.7*e* 23), but not during wake-like regimes (B; *t* = −0.6, *p* = 0.58). The PLI is consistently large in sleep-like dynamics (G-H), unlike in wake-like dynamics where the PLI is near zero (E-F), suggesting systematic phase relations between brain regions in synchronous states consistent with travelling slow waves experimentally observed during sleep. Likewise, the PLI of excitatory and inhibitory populations is significantly different in sleep-like (H; Independent Student’s T-test, *t* = −9.5, *p* = 4.2*e* 21), but not wake-like (F; *t* = −0.5, *p* = 0.60) states, altogether possibly suggesting a previously unidentified role of inhibition in the emergence of long-range synchrony in sleep-like activity.

### 2.4 Responsiveness to external stimulation

After reproducing features of spontaneous dynamics between brain states, we test the hypothesis that changing adaptation in the TVB-AdEx model can also explain differences in empirically observed stimulus-evoked brain responses, with stimuli encoded in more sustained, widespread, reliable, and complex patterns during conscious states [3]. To this end, a square wave of 0.1*Hz*, matching the magnitude of stochastic drive, with 50*ms* duration was input to the firing rates of the transfer function for excitatory populations in the right premotor cortex of the TVB-AdEx simulation during awake-like and slow wave sleep-like conditions, as in previously published empirical studies [4].

Figure 5 illustrates the effect of perturbing the large-scale network defined by TVB-AdEx models.

**Figure 5.**
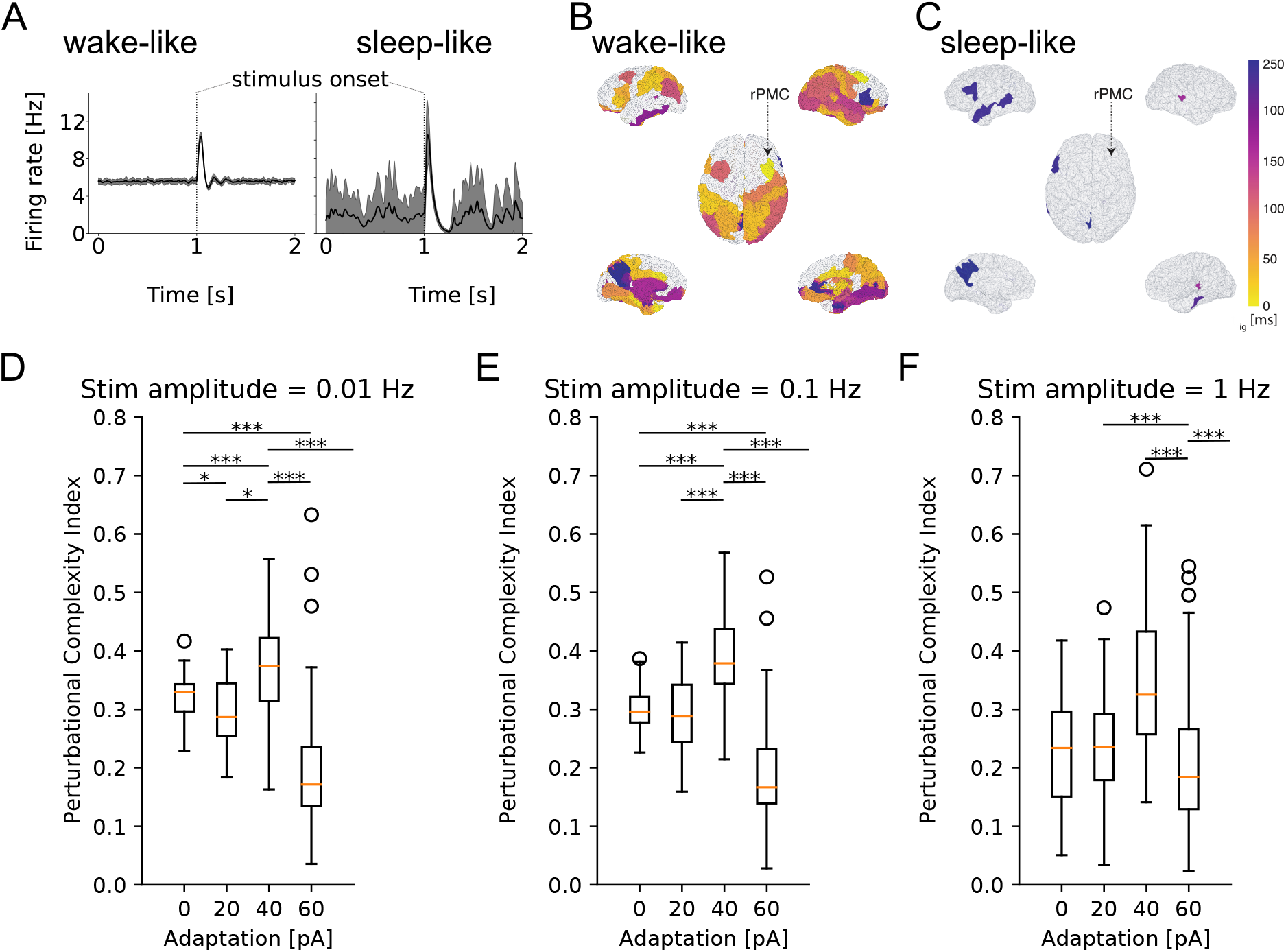
Increased responsiveness to perturbations in simulated wake-like compared to sleep-like states. Excitatory firing rate of a simulated brain region in time during wake-like (A) and sleep-like dynamics, in response to a stimulus. Black lines show the mean across 40 realizations, reminiscent of event-related potentials (ERPs), and grey shaded areas show the standard deviation from the mean across realizations. (B-C) Spatio-temporal propagation of responses to stimuli in wake-like (B) and sleep-like (C) states. Color plotted on the brain surface indicates earliest time at which each region becomes significantly different from its pre-stimulus baseline (see Methods), with earlier times in lighter colours. Regions in white do not present significant differences in firing rate in response to the stimulus. In wake-like states, stimulus responses recruit more brain regions and produce more complex spatio-temporal patterns, reminiscent of empirical observations. (D-F) Box plots of perturbational complexity index (PCI) measurements from 40 realizations of wake-like and sleep-like simulations with increasing stimulus amplitudes (panels left to right). Significant changes in the PCI are observed when the spike frequency adaptation (*b_e_*) is varied (one-way Kruskal-Wallis test; p<0.05 for each group of adaptation values, *b_e_* = 0, 20, 40, 60, for each stimulus value (0.01, 0.1, and 1 Hz). Results of post-hoc Conover test for multiple comparisons between values of *b_e_* are shown in the figure, where * denotes p<0.05, ** is p<0.01, and *** is p<0.001). In high-adaptation, sleep-like regimes, a sharp drop in PCI is observed, denoting more spatio-temporally complex responses in wake-like compared to sleep-like states, consistent with experiments [3, 4].

The effect of an external stimulus is apparent for both deep sleep-like and wake-like states (Fig. 5A). The average traces of the stimulated region are shown in black, and take into account the 40 realisations shown in grey. To examine the spread of activity following perturbations, the time at which the excitatory firing rate of each region becomes significantly different from the unstimulated baseline (prior to perturbation) is plotted using a color map showing earlier significant changes in brighter colors. Here, we find that responses are more widespread across time and space across brain regions in wake-like (Fig. 5B) than sleep-like dynamics (Fig. 5C), corresponding to experimental observations in response to Transcranial Magnetic Stimulation (TMS) [3].

To better characterize the spatiotemporal effects of stimuli, the perturbational complexity index (PCI), used in previous experimental works involving TMS, was computed (see Methods). A low PCI value indicates a “simple” response to stimulus, while a high PCI value indicates more “complex” response, typically propagating more effectively to different brain areas [4]. The models were perturbed with stimuli smaller (0.01*Hz*) than the stochastic drive, disguised within the noise (0.1*Hz*), and larger than the drive (1.0*Hz*) for simulations in which the value of spike frequency adaptation (*b_e_*) was varied between 0 pA and 60 pA. As shown in Fig. 5D-F, computing the PCI from from the TVB-AdEx model shows that PCI values are typically higher for lower-adaptation, wake-like regimes than for higher-adaptation, slow-wave sleep like regimes. In particular, a sharp drop in PCI values is observed between *b* = 40pA and *b* = 60 pA, suggesting an abrupt transition between highly responsive asynchronous and less responsive slow-wave dynamics as adaptation increases. For each value of noise, a one-way ANOVA revealed significant differences between PCI distributions across *b* values (*p* < 0.0001), with multiple comparisons highlighting that the PCI was significantly larger for wake-like than sleep-like conditions, in particular for lower-amplitude stimuli. The same behavior was observed when comparing awake subjects with subjects in slow-wave sleep [3, 4]. Also note the wider distribution of PCI values in sleep-like simulations, suggesting more variable responses for each realisation of the same stimulus and therefore less reliable stimulus encoding.

## 3 Discussion

In this paper, we demonstrated that biologically-informed scale-integrated mean-field models [9] can be used to simulate large-scale brain networks using the TVB platform in EBRAINS. The coupled mean-field models comprising the TVB-AdEx are derived from networks of AdEx neurons and display whole-brain asynchronous and slow-wave dynamics when wired following white matter tracts from a human connectome. These results demonstrate the natural emergence of empirically observed patterns of macroscopic brain dynamics from simulated changes at microscopic scales. The TVB-AdEx integration in EBRAINS is also of interest as EBRAINS human brain atlas services will be able to provide a large degree of cytoarchitectural detail such as region-specific neurotransmitter densities and cell types and densities and thus add to the biological realism of these virtual brain models. The vertical integration across scales is provided by TVB-AdEx-type models, taking advantage of the Big Data in EBRAINS.

TVB-AdEx mean-field models constituting each node of the connectome are designed by construction to approximate the mean and covariance of the firing rate in spiking neural networks exhibiting stable dynamics in asynchronous irregular regimes [20]. This model was extended to two neuronal populations, excitatory neurons with adaptation and inhibitory neurons [9], but this extension has limitations. Importantly, the model is imprecise when adaptation varies within a range larger than described here (for adaptation values higher than 100pA [9]) and when fast synchronous dynamics like oscillations in the gamma range (between 40 Hz and 80 Hz) [20], spindles, or ripples occur. This model is therefore likely not directly suitable for understanding the nuances associated with particular microscopic motifs comprised by transiently communicating assemblies that likely encode relevant neural information. The model is appropriate to the type of general dynamics presented here, wake-like AI and deep sleep-like slow-waves, describing large scale phenomena with relatively slow time scales. By smoothing microscopic details, we have built a computationally tractable bridge from microscopic to macroscopic scales, to elucidate how general dynamical phenomena relate to differences in neuronal interactions.

After integration in TVB, the resulting TVB-AdEx model displays a number of interesting features and several exciting perspectives for future work. A first result is the emergence of synchrony across brain regions in the presence of adaptation. In this case, the TVB-AdEx model displays synchronized slow waves with structured phase relations at a macroscopic, brain-wide level (Fig. 3). This is consistent with the synchronized slow-wave dynamics observed during deep sleep in the brain [1, 18, 19]. When the same model is set into the asychronous-irregular regime due to the loss of spike-frequency adaptation, as is the case in the presence of acetylcholine and other neuromodulators present in higher concentrations during waking states [29], the large-scale network displays a much lower level of synchrony (Fig. 3), consistent with asynchronous dynamics typically seen in awake and aroused states [1, 18, 19]. These different levels of synchrony are therefore emergent properties of the large-scale network.

A second main result is that evoked dynamics are also adaptation-dependent; When the network displays synchronized Up- and Down-states, a stimulus typically evokes a high amplitude, simple response that remains local in space and time. When the model is in the asynchronous regime, the same stimulus evokes responses that are weaker in amplitude, but that propagate in a more elaborate way through space and time. The PCI measure applied to these two states match the experimental observations [3, 4, 5, 6]. Again, this is an emergent property of the large-scale network.

What are possible mechanisms for such differences? A previous study [15] showed that in balanced networks, not all states are equal and that asynchronous states, despite their apparently noisy character, can display higher responsiveness and support propagation of stimuli. This enhanced responsiveness of AI states can be explained by the combined effects of depolarization, membrane potential fluctuations, and conductance state. It was proposed as a fundamental property to explain why the activity of the brain is systematically asynchronous in aroused states [15]. The present results are in full agreement with this mechanism, which manifests here in the asynchronous state as a propagation further in time and space, across many brain areas, associated with higher values of the PCI.

We believe that this work opens several perspectives. First, the enhanced propagation of perturbations during wake-like states could be used as a basis to explain why stimuli are perceived in asynchronous regimes, and what kind of modulation of the network activity could support phenomena such as attention and perception. Second, mean-field models can be set to also display pathological states, such as hyperactive or hypersynchronized states, and the TVB-AdEx model could be used to investigate seizure activity [30]. Other features, such as neuronal heterogeneity, are also beginning to be included in mean-field models, [31], paving the way for enhancing biological realism in future versions of TVB-AdEX models.

In TVB, connectivity depends on the intermediate spatial resolution of coarse-graining. Here, the brain was parcellated in 68 regions, with each mean-field representing a substantially large brain area. TVB allows such simple simulations using a few tens of nodes, taking into account the rough long-range connectivity according to the connectome resolved in tractography of human dMRI. TVB can also simulate much finer-grained connectivity, by defining a larger number of nodes (usually on the order of hundreds to hundreds of thousands, approaching the resolution of cortical columns) [32]. In such vertex-based simulations that shall follow the present work, local connectivity is determined by intracortical connections, whereas the white-matter connectome from dMRI used here captures effects of longer-range cortico-cortical fibers.

Early stimulation studies in humans and in particular in rodents are pioneering the use of high-resolution simulations, demonstrating subtler influences of the connectome in scaffolding signal propagation through brain networks [33, 34]. In those studies, network nodes were equipped with generic neural mass dynamics, Andronov-Hopf-oscillators, which are theoretically appealing for their mathematical simplicity, but are limited with regard to biophysical interpretability of the results. The inclusion of high-resolution data from tracer studies in the Allen Institute was recently demonstrated in virtual mouse brain models to significantly increase the predictive power [35]. As well, the inclusion of subject-specific, personalized connectomes in virtual brain models significantly outperforms generic simulations in predictive inter-individual variability [35, 36]. These studies point together to the importance of personalized brain network models in future clinical applications and affords novel methods supporting such goals [37, 38]. The virtual brains in Spiegler et al [33, 34] captured the emergence of well-known resting state networks known during spontaneous activity, but also functionally specific brain responses to stimulation of regions along the processing chains of sensory systems from periphery up to primary sensory cortical areas. The latter responses heavily relied on the Default Mode Network (DMN) and were suggestive of the DMN playing a mechanistic role between functional networks. But neither brain state dependence, nor biological interpretation of the neural mass model parameters was possible, as it requires the incorporation of biological complexity and integration across scales provided here by the here by the TVB-AdEx approach. Ongoing efforts in EBRAINS aim to enrich high-resolution brain models with detailed information on regionally-variant physiological features (neurotransmitters, receptor densities, cell types, and densities) to next build the Virtual Big Brain, a high-resolution multi-scale brain network, which will be continuously updated and available to the community. The drawback of such fine grained simulations is that they typically require large computing resources as provided by EBRAINS, while coarse grained TVB simulations, as presented here, can easily be run on a standard workstation. To summarize, for the sake of the initial release of the TVB-AdEx models, we offer a relatively coarse parcellation, which will become more refined and personalized in future work.

Finally, it is interesting to note that the global properties used here to characterize neural dynamics across brain states - synchrony, frequency spectra, responsiveness, and PCI - all reflect neural correlates of consciousness [39, 40, 41, 42]. It has even been argued that asynchronous dynamics (so-called ‘activated EEG’) is so far one of the most “sensitive and reliable” neural correlate of consciousness [39]. This further emphasizes the promise of the present modeling approach to understanding dynamics associated with conscious and non-conscious states, with broad potential applications in medicine and computation.

## Acknowledgments

This research was funded by the European Community (Human Brain Project, H2020-945539), the Centre National de la Recherche Scientifique (CNRS, France), the ANR PARADOX, and the ICODE excellence network. The authors would like to thank Nuria Tort-Colet, Matteo di Volo, Mallory Carlu, Michele Farisco, Thierry Nieus, and Mavi Sanchez-Vives for helpful feedback on the work in progress. Many thanks to EBRAINS for help with implementation and to Kevin Ancourt for wrangling the Git repository.

## Code availability

A python-based open-access code to run the present whole-brain model will be accessible online in the EBRAINS platform (https://ebrains.eu) as a companion to the publication of the present article. The scripts are also accessible on Github at https://gitlab.ebrains.eu/kancourt/tvb-adex-showcase3-git.

## Competing interests statement

The authors declare no competing interest.

JG and LK wrote the models, JG, LK, BHY, and DD characterized the models and ran simulations. TAEN, LK, and JG performed the analyses. JG, TAEN, VJ, and AD designed the work and wrote the manuscript. AD and VJ supervised the work.

## 4 Materials and Methods

Three types of models are used in this work: a network of spiking neurons, a mean-field model of this network, and a network of mean-field models implemented in The Virtual Brain (TVB). Here we describe these models successively.

### 4.1 Spiking network model

We considered networks of integrate-and-fire neuron models displaying spike-frequency adaptation, based on two previous papers [14, 8]. We used the Adaptive Exponential (AdEx) integrate-and-fire model [21]. We considered a population of *N* = 10^4^ neurons randomly connected with a connection probability of *p* = 5%. We considered excitatory and inhibitory neurons, with 20% inhibitory neurons. The AdEx model permits to define two cell types, “regular-spiking” (RS) excitatory cells, displaying spike-frequency adaptation, and “fast spiking” (FS) inhibitory cells, with no adaptation. The dynamics of these neurons is given by the following equations:

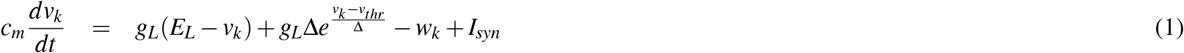

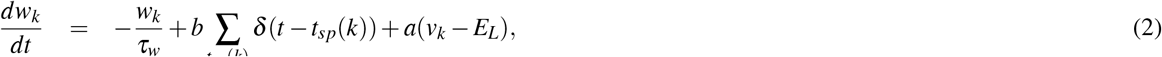

where *c_m_* = 200 pF is the membrane capacitance, *v_k_* is the voltage of neuron *k* and, whenever *v_k_ > v_peak_* = −47.5 mV for inhibitory neurons and *v_k_ > v_peak_* = −40.0 mV for excitatory at time *t_sp_*(*k*), *v_k_* is reset to the resting voltage *v_reset_* = −65 mV and fixed to that value for a refractory time *T_refr_* = 5 ms. The voltage threshold *v_thr_* is −50 mV. The leak term *g_L_* had a fixed conductance of *g_L_* = 10 nS and the leakage reversal *E_L_* was of 65 mV for inhibitory and 63 for excitatory. The exponential term had a different strength for RS and FS cells, i.e. Δ = 2mV (Δ = 0.5mV) for excitatory (inhibitory) cells. Inhibitory neurons were modeled as fast spiking FS neurons with no adaptation (*a* = *b* = 0 for all inhibitory neurons) while excitatory regular spiking RS neurons had a lower level of excitability due to the presence of adaptation (while *b* varied in our simulations we fixed subthreshold adaptation *a* = 0 nS and *τ_w_* = 500 ms).

The synaptic current *I_syn_* received by neuron *i* is the result of the spiking activity of all neurons *j* ∈ pre(*i*) pre-synaptic to neuron *i*. This current can be decomposed in the synaptic conductances evoked by excitatory E and inhibitory I pre-synaptic spikes

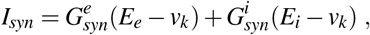

where *E_e_* = 0mV (*E_i_* = −80mV) is the excitatory (inhibitory) reversal potential. Excitatory synaptic conductances were modeled by a decaying exponential function that sharply increases by a fixed amount *Q_E_* at each pre-synaptic spike, i.e.:

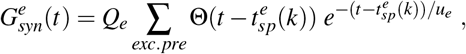

where Θ is the Heaviside function, *u_e_* = *u_i_* = 5ms is the characteristic decay time of excitatory and inhibitory synaptic conductances, and *Q_e_* = 1.5 nS (*Q_i_* = 5 nS) the excitatory (inhibitory) quantal conductance. Inhibitory synaptic conductances are modeled using the same equation with *e* → *i*. This network displays two different states according to the level of adaptation, *b* = 0 pA for asynchronous irregular states, and *b* = 60 pA for Up-Down states (see [8] for details).

### 4.2 Mean-field models

We considered a population model of a network of AdEx neurons, using a Master Equation formalism originally developed for balanced networks of integrate-and-fire neurons [20]. This model was adapted to AdEx networks of RS and FS neurons [8], and later modified to include adaptation [9]. The latter version is used here, which corresponds to the following equations using Einstein’s index summation convention where sum signs are omitted and repeated indices are summed over:

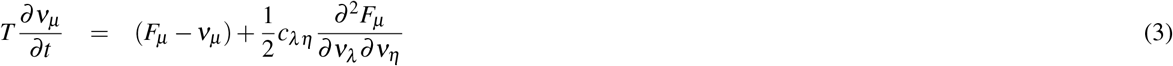

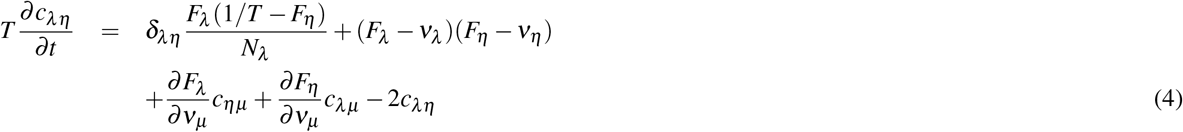

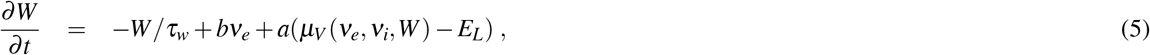

where *μ* = {*e, i*} is the population index (excitatory or inhibitory), *ν_μ_* the population firing rate and *c_λη_* the covariance between populations *λ* and *η*. *W* is a population adaptation variable [9]. The function *F_μ_*_={*e,i*}_ = *F_μ_*_={*e,i*}_(*ν_e_, ν_i_,W*) is the transfer function which describes the firing rate of population *μ* as a function of excitatory and inhibitory inputs (with rates *ν_e_* and *ν_i_*) and adaptation level *W*. These functions were estimated previously for RS and FS cells and in the presence of adaptation [9].

At the first order, i.e. neglecting the dynamics of the covariance terms *c_λη_*, this model can be written simply as:

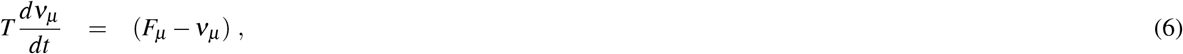

together with Eq. 5. This system is equivalent to the well-known Wilson-Cowan model [43], with the specificity that the functions *F* need to be obtained according to the specific single neuron model under consideration. These functions were obtained previously for AdEx models of RS and FS cells [8, 9] and the same are used here.

For a cortical volume modeled as a two populations of excitatory and inhibitory neurons, the equations can be written as:

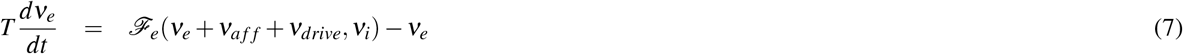

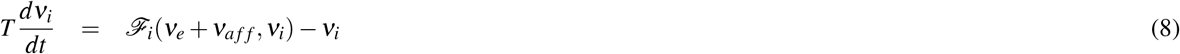

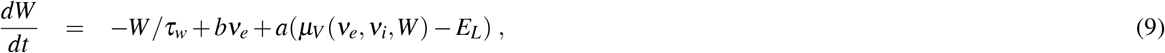

where *ν_aff_* is the afferent thalamic input to the population of excitatory and inhibitory neurons and *ν_drive_* is an external noisy drive simulated by an Ornstein-Uhlenbeck process. The function *μ_V_* is the average membrane potential of the population and is given by

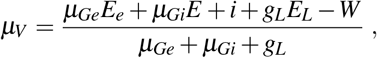

where the mean excitatory conductance is *μ_Ge_* = *ν_e_K_e_u_e_Q_e_* and similarly for inhibition.

This system describes the population dynamics of a single region, and was shown to closely match the dynamics of the spiking network [9].

### 4.3 Networks of mean-field models

Extending our previous work at the mesoscale [9, 44] to model large brain regions, we define networks of mean-field models, representing interconnected brain regions (each described by a mean-field model). We considered interactions between cortical regions as excitatory, while inhibitory connections remain local to each region. The equations of such a network, expanding the two-population mean-field (Eq. 7), are given by:

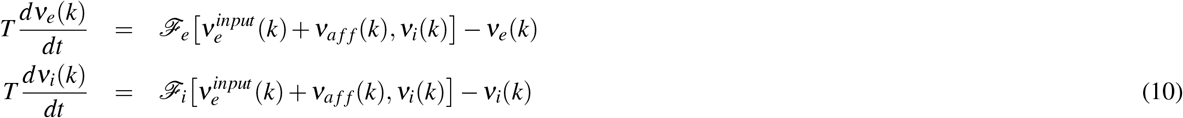

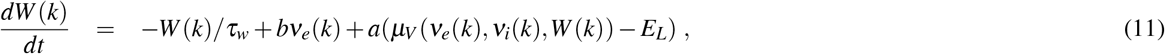

where *ν_e_*(*k*) and *ν_i_*(*k*) are the excitatory and inhibitory population firing rates at site *k*, respectively, *W* (*k*) the level of adapation of the population, and 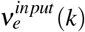 is the excitatory synaptic input. The latter is given by:

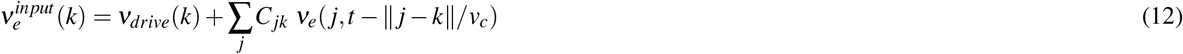

where the sum runs over all nodes *j* sending excitatory connections to node *k*, and *C_jk_* is the strength of the connection from *j* to *k* (and is equal to 1 for *j* = *k*). Note that *ν_e_*( *j, t − ||j − k||/v_c_*) is the activity of the excitatory population at node *k* at time *t − ||j − k||/v_c_* to account for the delay of axonal propagation. Here, ||*j − k*|| is the distance between nodes *j* and *k* and *v_c_* is the axonal propagation speed.

### 4.4 Spontaneous activity

The Phase-Lag Index (PLI) was computed for each pair of nodes, averaged over simulation time. The Hilbert transform is employed to extract the phase *ψ*(*t*) of the time series. From there, the PLI, given by

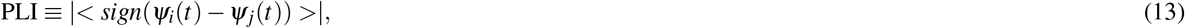

is computed for nodes *i* and *j*, where < · > denotes averaging over time [45]. One may note that the PLI takes values between 0 (random phase relations or perfect synchrony) and 1 (perfect phase locking). In this work we report the mean PLI over all time epochs for excitatory and inhibitory firing rates of each region pair for each adaptation value.

#### 4.4.1 Evoked activity

Here, we compute the Perturbational Complexity Index (PCI) in response to a localized square wave stimulus, over the firing rates of all nodes of a TVB-AdEx simulation, following the method proposed by Casali *et al* [4]. This is done for multiple trials with the same stimulus delivered to the same node at a random point in time and with different realizations of noise. The PCI is the ratio of two quantities: the Lempel-Ziv algorithmic complexity and the source entropy [4]. To compute both quantities, firing rates *ν*(*t*) must be binarized to produce significance vectors *s*(*t*). First, the trials are aligned to stimulation time, considering only the 300*ms* before and after stimulus onset. Then, each node’s firing rate is re-scaled and mean and standard deviation given by pre-stimulus activity averaged over nodes. Afterwards, all pre-stimulus firing rates are randomized across time bins, this procedure being repeated 500 times. The threshold for significance *T* is then given by the one-tail percentile of the maximum absolute value over all repetitions within a series of 20 trials. For each trial of those 20 trials, we can then write *s*(*t*) = 1 whenever post-stimulus *ν*(*t*) *> T* and *S*(*t*) = 0 otherwise. For what follows, we concatenate all *s*(*t*) vectors from all simulation nodes into one single significance vector *S*(*t*) per trial.

The Lempel-Ziv complexity *LZ*(*S*) is the length of the ‘zipped’ vector *S*(*t*), i.e. the number of possible binary ‘words’ that make up the binary vector *S*(*t*). Briefly, *S*(*t*) is sectioned successively into consecutive words of between one and *N_t_* characters where *N_t_* is the total length of *S*(*t*). Scanning sequentially through all words, each new encountered word is added to a ‘dictionary’, and *LZ*(*S*) is the total number of words in the dictionary at the end of the procedure.

The spatial source entropy *H*(*S*) is given by:

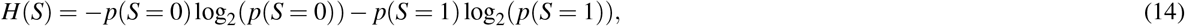

where log_2_ denotes the base-two logarithm.

The PCI can then be expressed as 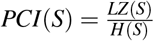.

